## Supplemental figures for "Non-synonymous to synonymous substitutions suggest that orthologs tend to keep their functions, while paralogs are a source of functional novelty"

### Supplementary figures

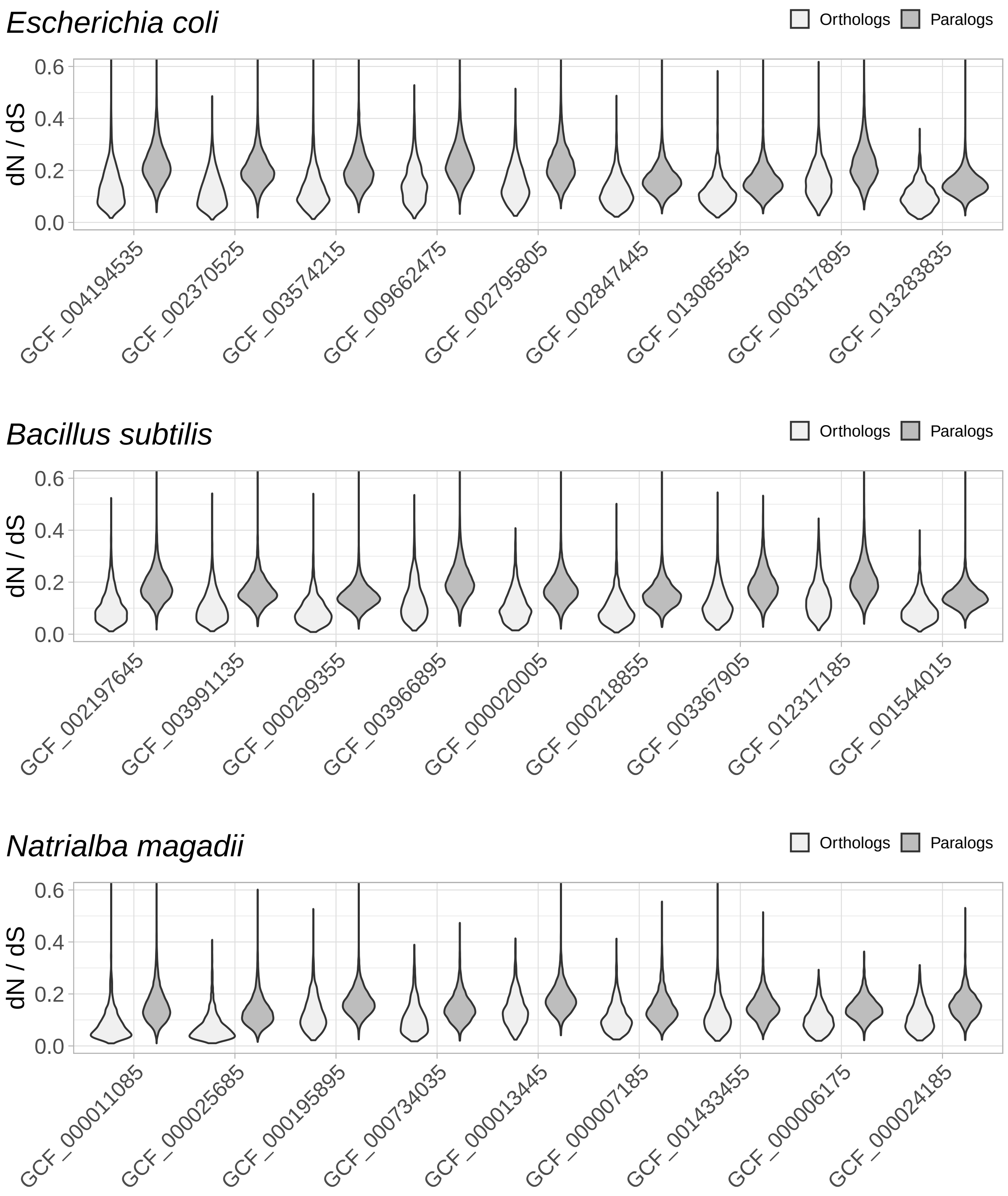


Figure S1: Full $dN/dS$ results obtained with InParanoid as a working definition of orthology.

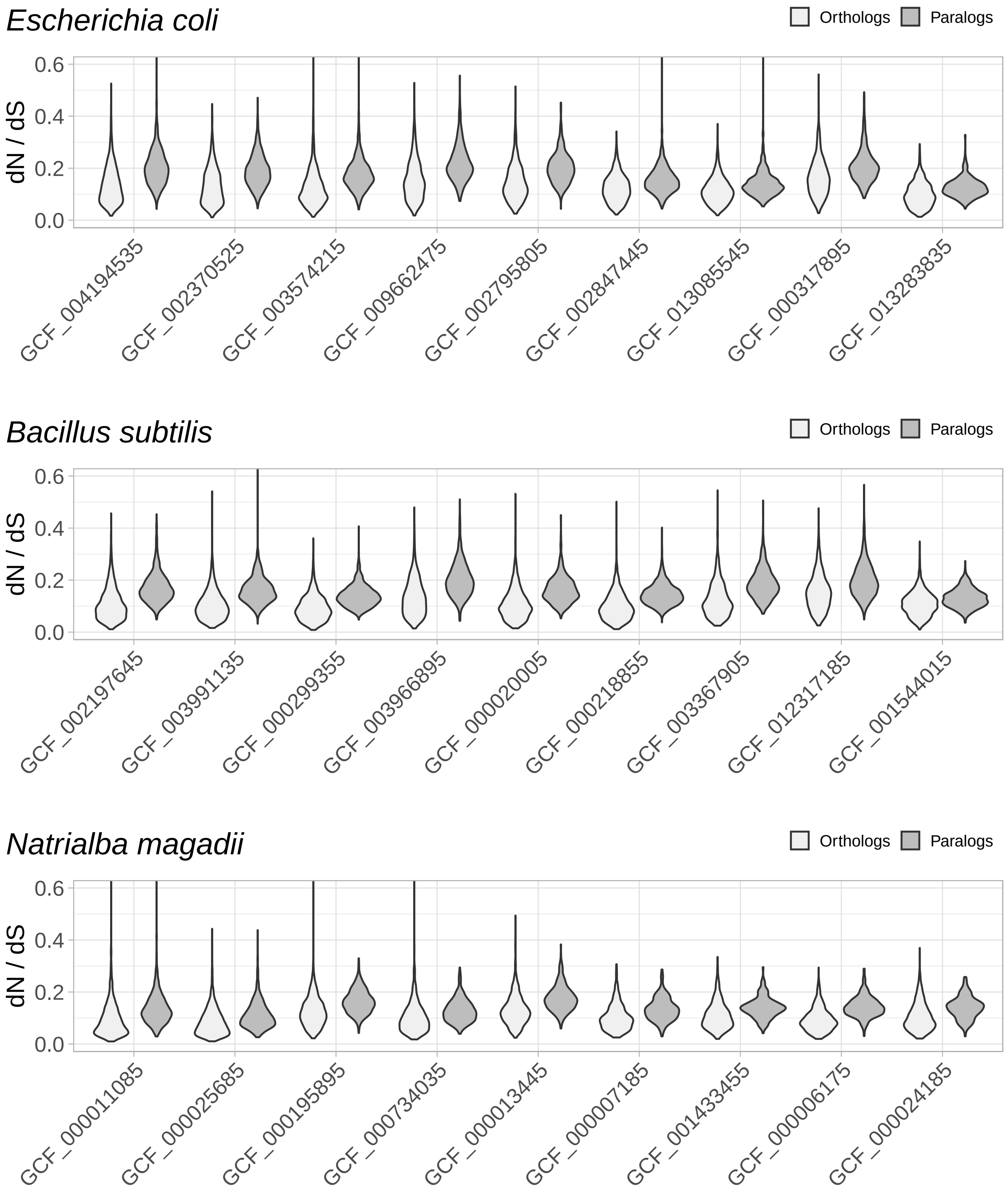


Figure S2: Full $dN/dS$ results obtained with the Orthologous Matrix (OMA) working definition of orthology.

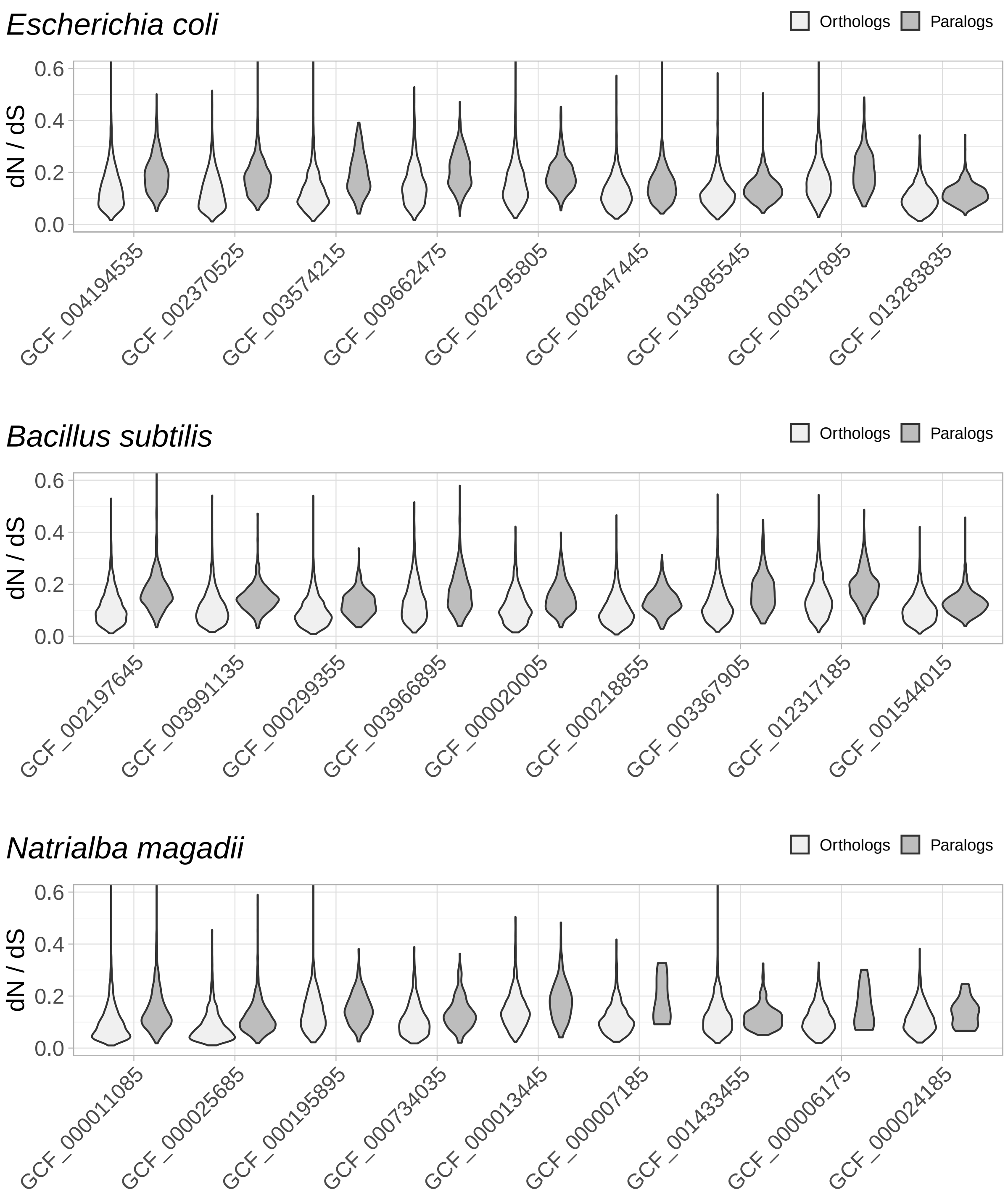


Figure S3: Full $dN/dS$ results obtained with the OrthoFinder working definition of orthology.

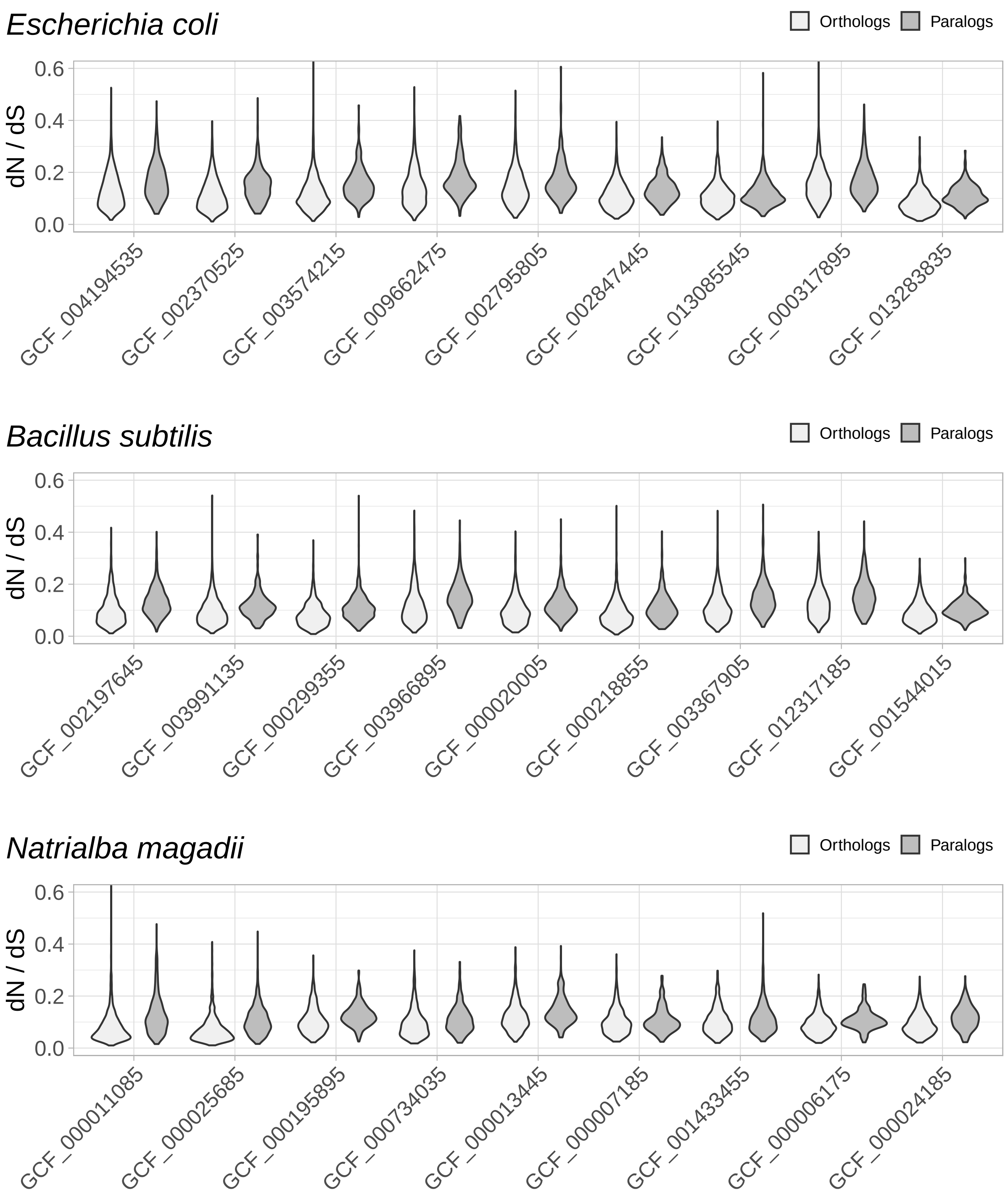


Figure S4: Full $dN/dS$ results obtained with the ProteinOrtho working definition of orthology.

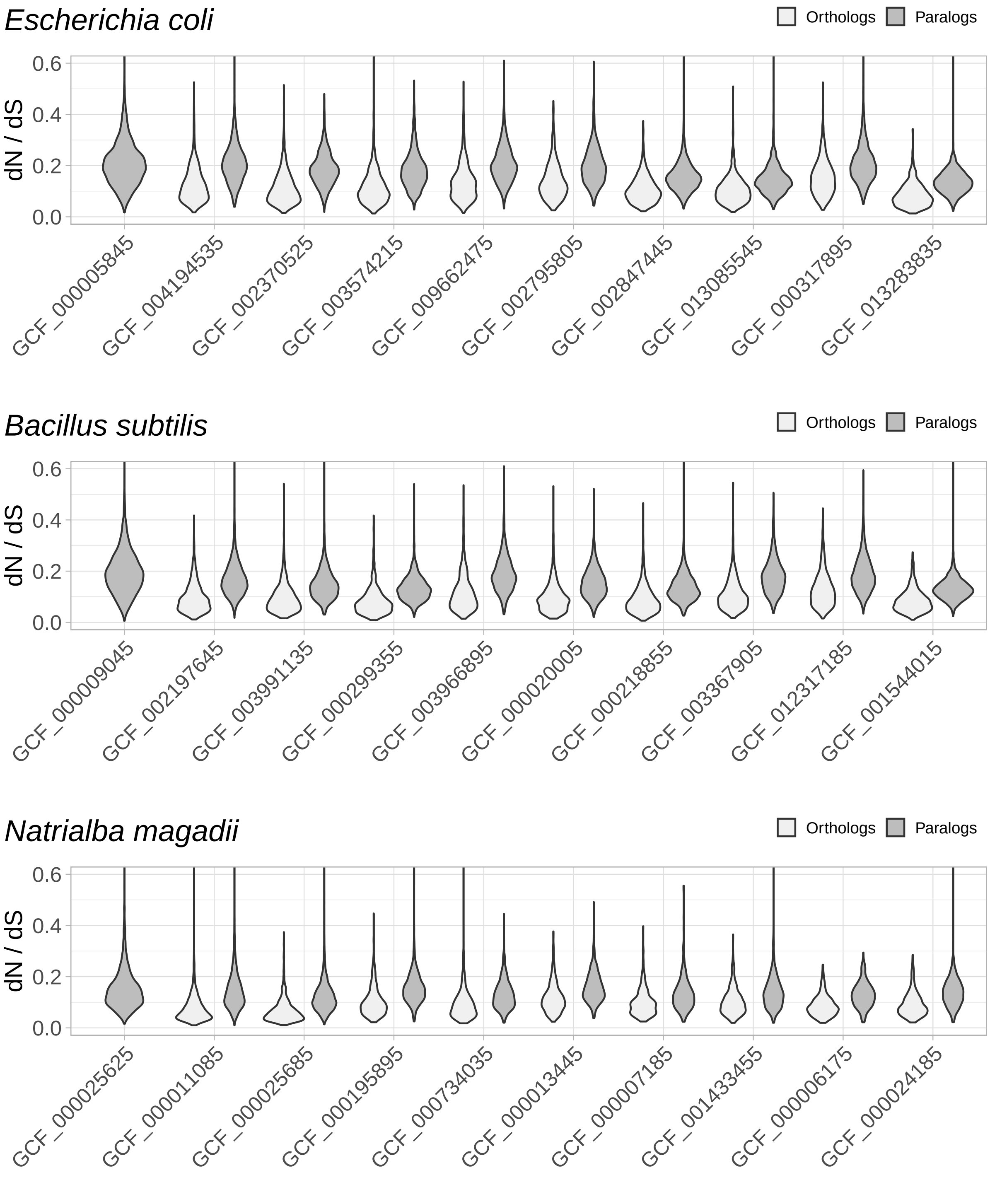


Figure S5: Full $dN/dS$ results obtained with alignments covering at least 80% of both proteins.

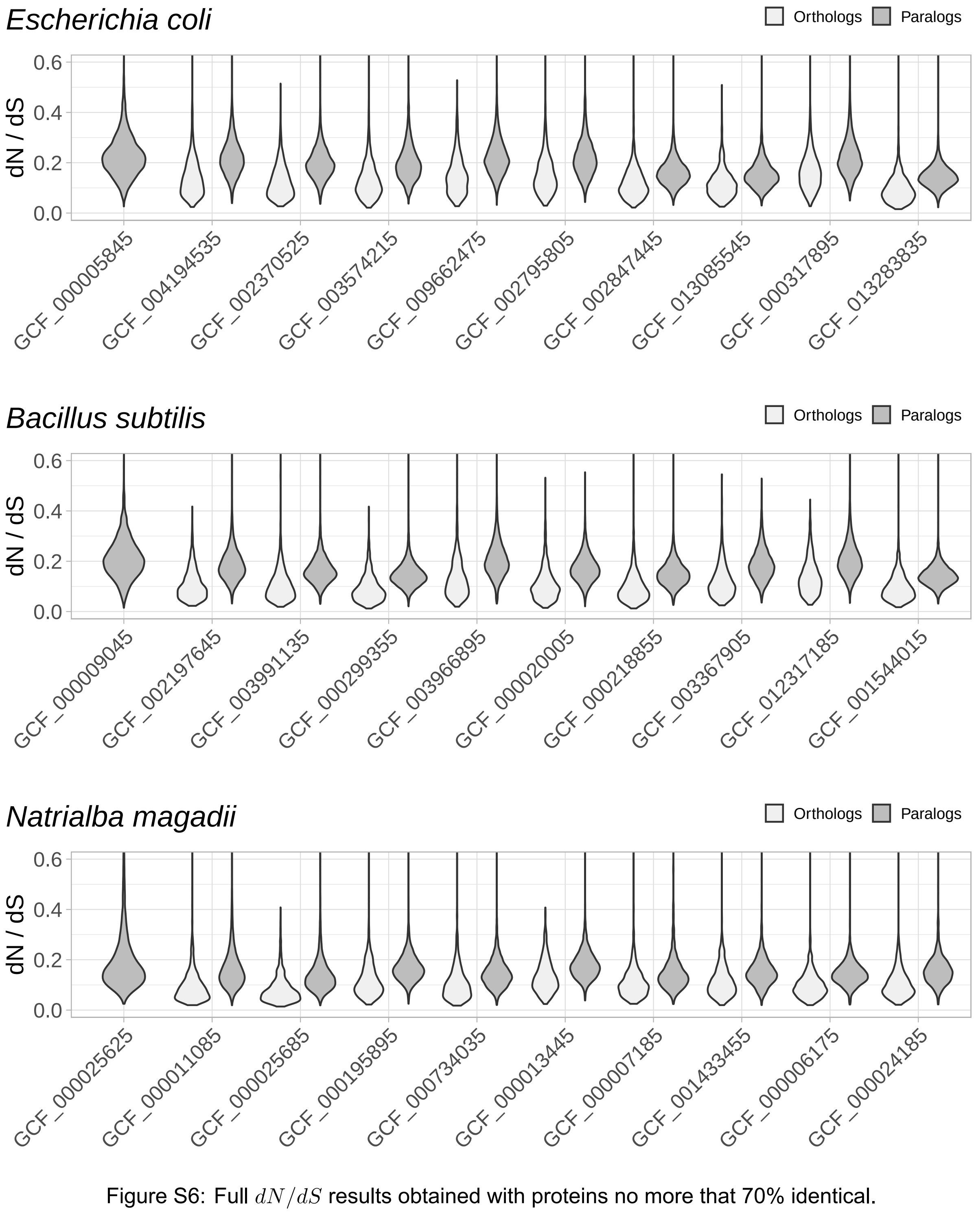


Figure S6: Full $dN/dS$ results obtained with proteins no more that 70% identical.

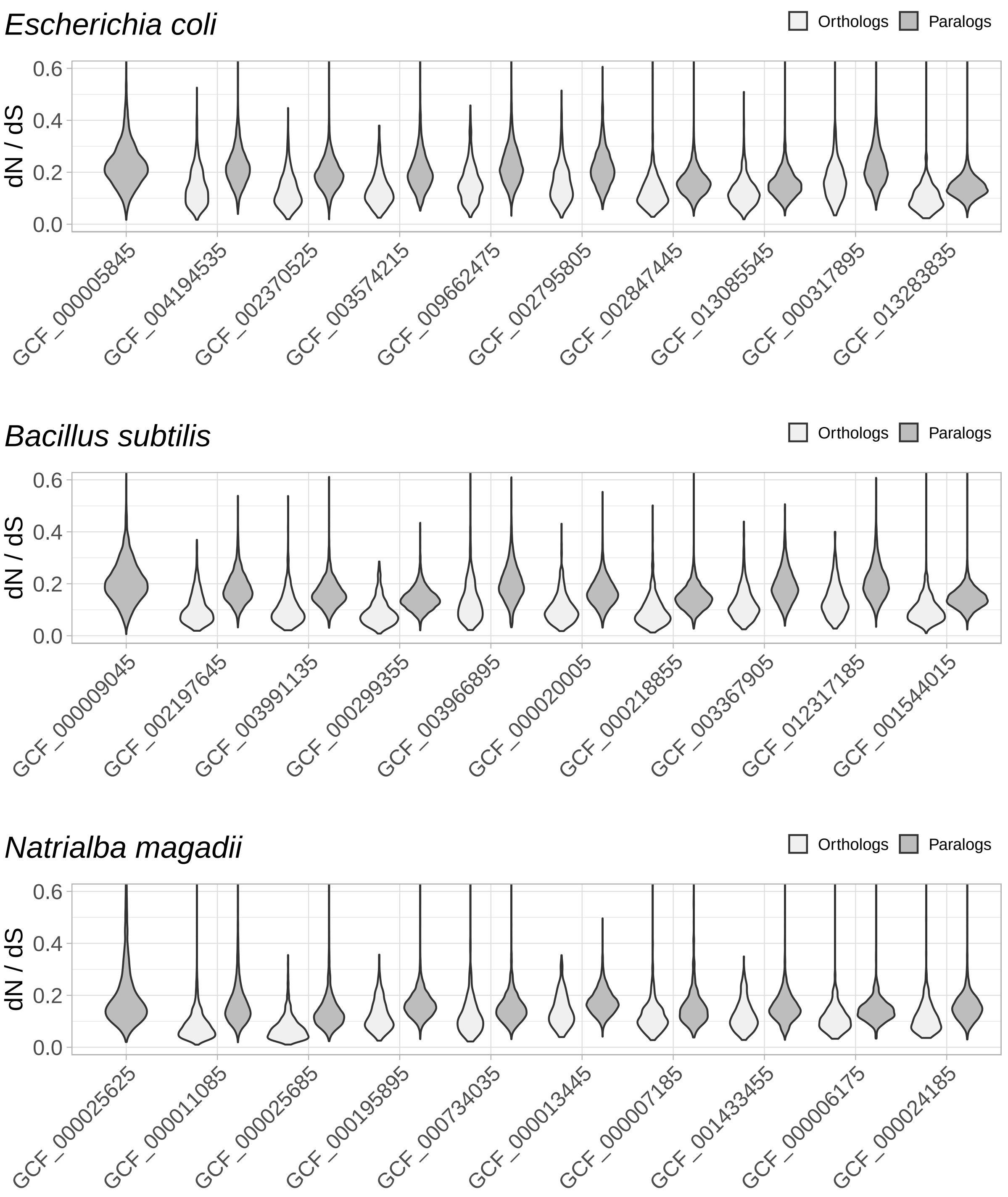


Figure S7: Full $dN/dS$ results obtained with proteins within usual codon usage.

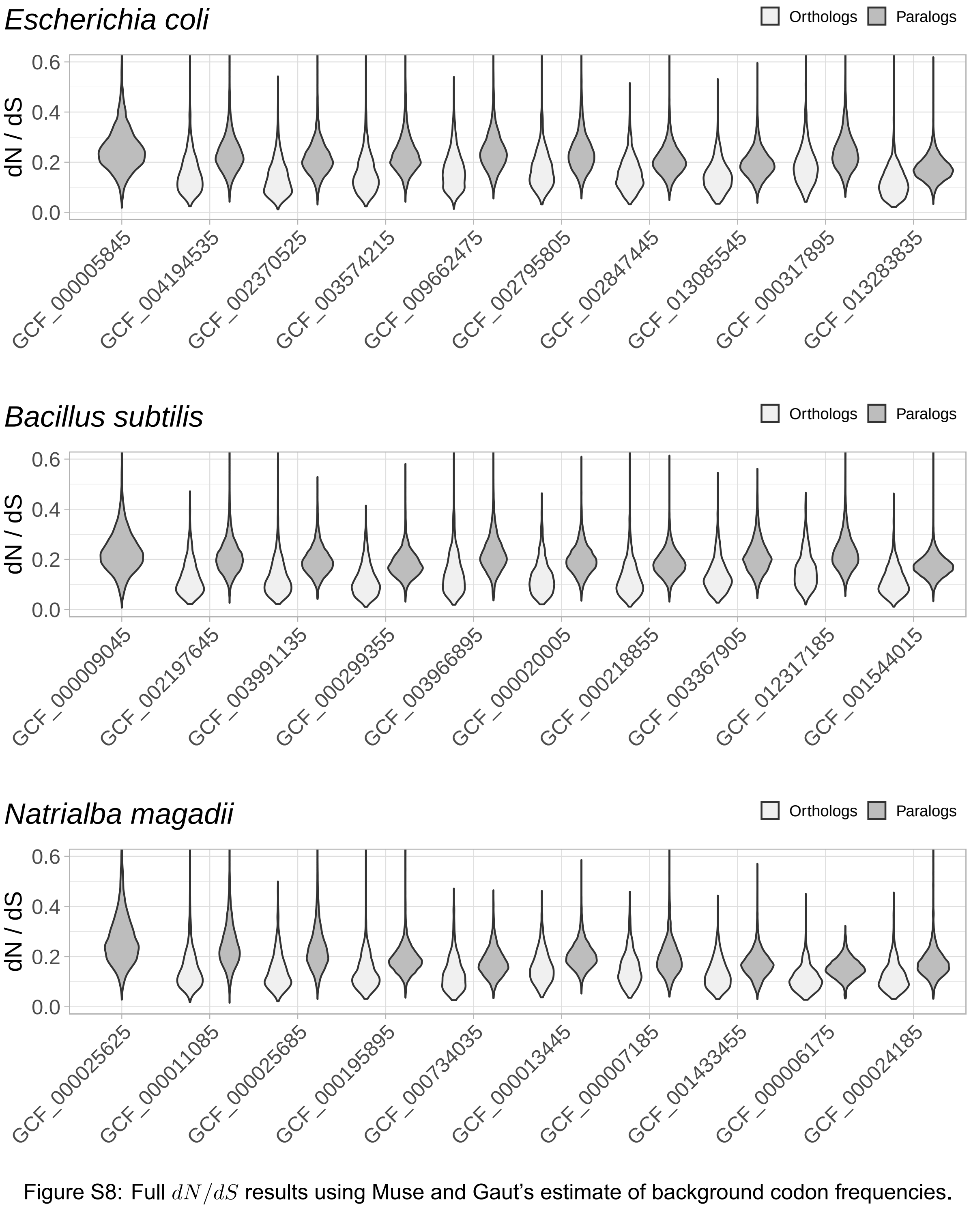


Figure S8: Full $dN/dS$ results using Muse and Gaut's estimate of background codon frequencies.
